# Sphere-sequencing unveils local tissue microenvironments at single cell resolution

**DOI:** 10.1101/2022.10.31.514509

**Authors:** Kristina Handler, Karsten Bach, Costanza Borrelli, Xenia Ficht, Ilhan E. Acar, Andreas E. Moor

## Abstract

The spatial organization of cells within tissues is tightly linked to their biological function. Yet, methods to probe the entire transcriptome of multiple native tissue microenvironments at single cell resolution are lacking. Here, we introduce sphere-sequencing, a method that enables the transcriptomic characterization of single cells within spatially distinct tissue niches. Sphere-sequencing of the mouse metastatic liver revealed previously uncharacterized zonated genes and ligand-receptor interactions enriched in different hepatic microenvironments and the metastatic niche.

## Introduction

The last years have seen rapid technological developments to capture both transcriptomic features as well as corresponding tissue coordinates. However, the resulting spatial resolution of most unsupervised array-based techniques precludes single cell analyses, while imaging-based techniques require prior knowledge for targeted panel assembly^1^. Therefore, existing methods cannot simultaneously probe the entire transcriptome of multiple tissue microenvironments at single cell resolution. Here, we introduce sphere-sequencing (sphere-seq), a novel method to enable the transcriptomic characterization of single cells within spatially distinct tissue niches.

Previously reported comparable methods, such as Paired-cell sequencing^2^, PIC-seq^3^, and Clump-seq^4^, analyze spatial communities of 2-10 cells together in bulk and therefore lack single cell resolution. Computational deconvolution can be applied to such datasets to approximate single cell transcriptomes, but this process is inherently imprecise, especially for genes that are co-expressed in multiple cells. Sphere-seq refines these approaches and achieves single cell resolution while simultaneously capturing larger communities of cells that reflect biologically relevant tissue microenvironments. Importantly, this enabled us to predict ligand-receptor (L-R) interactions which are not only significantly enriched in a single cell RNA sequencing (scRNA-seq) dataset, but actually spatially co-occur in specific microenvironments.

Sphere-seq is based on partial tissue dissociation followed by size-dependent sorting of individual Ø 200-450 μm cellular communities (spheres) with a large fragment biosorter into a 96 well plate; one sphere per well. Subsequently, spheres are dissociated into single cells within each well and hashed with a sphere-specific lipid-tagging barcode^5^ to preserve information about a cell’s neighborhood prior to pooling for scRNA-seq (Fig. 1a). A species-mixing experiment indicated that 95% of cells are correctly assigned to their sphere of origin, confirming the validity of our approach (Supplementary Fig. 1a-d). We used sphere-seq to dissect spatially distinct niches in the metastasis-bearing mouse liver. The liver is characterized by a spatially repetitive organization into hexagonal hepatic lobules, in which blood flows from portal to central veins through sinusoids flanked by linear cords of hepatocytes (Fig. 1b). Differential oxygen and nutrient availability along the central-portal axis imprints zonation of cell functions and gene expression patterns in liver resident cells, particularly in hepatocytes and liver endothelial cells (LECs)^2,6–8^. Furthermore, multiple studies have reported zonation of immune cells, including liver-resident macrophages called Kupffer cells (KCs), T and NKT cells, which were all found to be periportally enriched^9–10^. However, the exact mechanisms of immune zonation and how it affects inflammatory responses or metastatic seeding are still to be investigated^9–10^.

## Results

### Sphere-sequencing reveals previously uncharacterized gene zonation in the metastasis-bearing mouse liver

We applied sphere-seq to mouse liver samples harvested two weeks after intrasplenic injection of colon cancer organoids (Supplementary Fig. 2a-d). Resulting single cell transcriptomes were annotated using known marker genes from the liver cell atlas^10^, assigned to their sphere of origin, and different cell type proportions per sphere were assessed (Fig. 1c,d, Supplementary Fig. 3a-c). We used a previously identified LEC zonation-specific gene expression signature^2^ to reconstruct the spatial origin of each sphere along the central-portal axis (lobule layers L1-L10). The accuracy of the reconstruction was corroborated by zonated gene expression in hepatocytes from pericentral and periportal spheres which matched previously published landmark genes^8^ (Supplementary Fig. 5a). This spatial ordering allowed us to characterize genes in LECs that were not reported as zonated in the reference dataset (Fig. 1e). Some of the identified zonated genes could be verified in healthy mouse liver and are therefore likely involved in homeostatic processes (Fig. 1f; Supplementary Fig. 5b). For example *Plpp1*, a phospholipid phosphatase, is centrally zonated, implying a link to pericentral lipogenesis^6^ while *Galnt15*, putatively involved in O-linked oligosaccharide biosynthesis, is portally zonated, which is in accordance with periportal gluconeogenesis^6^.

Next, we designed a 100 gene panel for highly-multiplexed fluorescence in situ hybridization (FISH, Molecular Cartography) which included marker genes for key cell types, as well as genes that displayed a spatially differentiated expression in sphere-seq (Supplementary Fig. 4a-d). This validated zonation patterns of metabolic genes in LECs (Fig. 1e,g; Supplementary Fig. 5c). Sphere-seq furthermore permitted the investigation of zonated genes in other cell types after assigning a sphere to periportal or pericentral zones according to LEC gene expression patterns. For example, *Vcam1* was found to be enriched in periportal KCs, which could be independently validated by Molecular Cartography (Fig. 1h). Interestingly, we did not find *Vcam1* to be significantly zonated in KCs in healthy livers suggesting its upregulation is a response to the metastatic process (Supplementary Fig. 5d). Upregulation of VCAM-1 during metastatic disease has been previously reported for LECs and may contribute to immune cell recruitment by interactions with its binding partner integrin α4β1^11^. To date, the function of VCAM-1 in KCs has not been well studied, but Okada et al. have described that VCAM-1 can mediate interactions of KCs and lymphocytes, which in turn promotes KC activation^12^.

This spurred us to perform L-R interaction analysis^13^ for KCs and lymphocytes (T and B cells) in periportal and pericentral zones. This analysis uncovered an enrichment of KC-expressed ligand *Vcam1* interacting with *α9β1, α4β7* and *α4β1* integrin complex receptors on T and B cells in periportal compared to pericentral zones (Fig. 1i, Supplementary Fig. 5e). Molecular Cartography could validate an enriched periportal gene expression of *Vcam1* and *Itgb1* (Integrin subunit β1) (Fig. 1j). A primary function of α4β7 and α4β1 integrins on T cells is mediating tissue homing and adhesion to the vessel wall^14^, indicating that periportal KCs may have upregulated *Vcam1* expression to promote lymphocyte recruitment. This is in line with an increase in predicted *Ccl3|Ccr5* interactions between periportal KCs and T cells, as CCR5 was reported to mediate the recruitment of T cells to the liver during acute inflammation^14^.

**Fig. 1:**
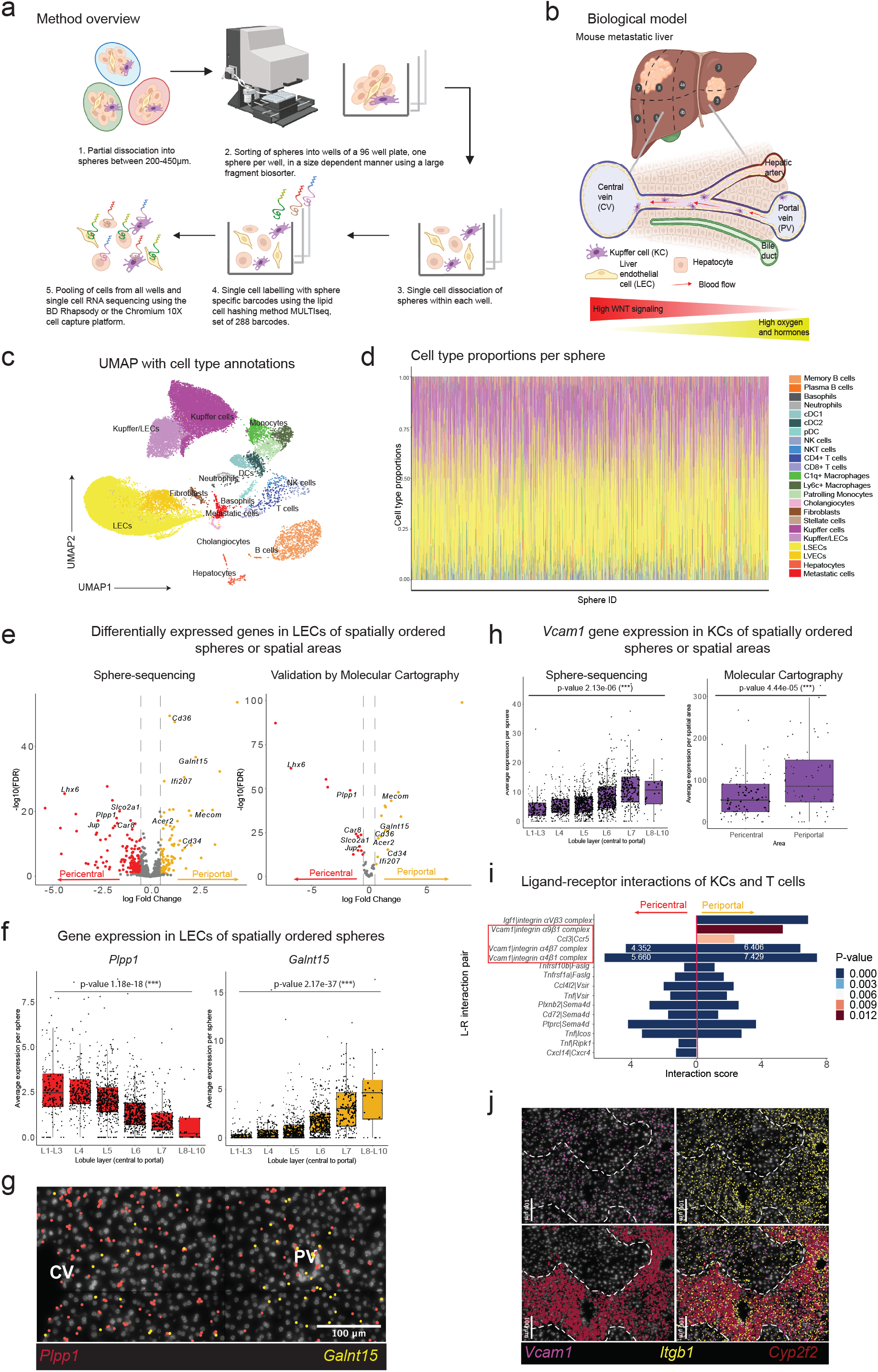
Overview of sphere-sequencing and its application to gene zonation in the metastasis-bearing mouse liver. **a**, Schematic illustration of sphere-sequencing workflow. **b**, Schematic drawing of liver macro- and microanatomy. **c**, Uniform manifold approximation and projection (UMAP) visualization of integrated samples of liver tissues (n=10: 9 injected with colon cancer organoids, 1 untreated) after filtering out spheres with less than 5 cells. Cells are clustered, annotated and coloured by their cell type. **d**, Barplot showing cell type proportions per sphere; only spheres with at least 5 cells are considered (n=1568 spheres across 10 samples). **e**, Differentially expressed genes (DEGs) of liver endothelial cells (LECs) in spatially ordered spheres (pericentral or periportal) displayed in volcano plots. Left, sphere-seq (1384 spheres across 9 samples). Right, validation with Molecular Cartography (155 areas across 4 samples). Coloured dots represent significantly enriched genes; red, enriched in pericentral zones; yellow, enriched in periportal zones. Gene labels indicate genes significantly enriched in both analyses. **f**, Boxplots showing gene expression in LECs of spatially ordered spheres (n= L1-L3: 137, L4: 214, L5: 402, L6: 409, L7: 196, L8-L10: 26 spheres across 9 samples). Black dots represent individual spheres. **g**, Molecular Cartography of *Plpp1* in red and *Galnt15* in yellow shown as an overlay with DAPI signal (white). CV: central vein; PV:portal vein. **h**, Boxplots of *Vcam1* gene expression in Kupffer cells (KCs) of spatially ordered spheres or spatial areas. Left, sphere-seq (like in f) (n= L1-L3: 115, L4: 206, L5: 379, L6: 387, L7: 182, L8-L10: 26 spheres across 9 samples); right, Molecular Cartography comparing pericentral and periportal (pericentral n=: 89; periportal n=66 areas across 4 samples). **i**, Predicted ligand-receptor (L-R) interactions between KCs and T cells in pericentral or periportal zones (n=9 samples). Interaction scores were calculated from sphere-seq data by CellPhoneDB, which uses a permutation test to generate p-values indicating significantly enriched L-R interactions. Interactions referenced in the main text are highlighted with red squares and white numbers indicate interaction scores. **j**, Representative Molecular Cartography of *Vcam1* (purple), *Itgb1* (yellow) and *Cyp2f2* (red) shown as an overlay with DAPI signal (white). For **e**,**f,** and **h** a negative binomial generalized log-linear model was used for statistical testing and p-values (Benjamini-Hochberg adjusted) of < 0.05 were considered significant (***<0.001, ** < 0.005, * < 0.05).

### Sphere-sequencing of metastases-bearing mouse liver reveals cell types and interactions specific to the metastatic-niche

Instead of grouping spheres according to liver zonation we next grouped them based on their distance to micro-metastatic sites. Spheres containing metastatic cells were defined as ‘proximal’ and spheres without metastatic cells as ‘distal’ to metastasis (Fig. 2a). First, we assessed cell-type proportions of key cell subsets and found that proximal areas had a significantly higher proportion of monocytes (Fig. 2b, Supplementary Fig. 6a). Segmentation and cell-type annotation of Molecular Cartography data validated those findings (Fig. 2c-d, Supplementary Fig. 6c), highlighting that sphere-seq faithfully recapitulates *in situ* cell-type proportions. We decided to investigate the monocyte subset more closely and found that proximal areas showed a trend for enrichment of *C1q+* macrophages (Fig. 2e). We could validate the increased abundance of *C1q+* monocytes within proximal areas in Molecular Cartography (Fig. 2f). *C1q+* macrophages are reportedly involved in T cell exhaustion and are an indicator of poor prognosis in many cancers^15^. Therefore we wanted to investigate cellular crosstalk between monocytes and T cells further. We found significantly more colocalisations between monocytes and T cells within proximal areas (Fig. 2g,h) suggesting a potential role for monocyte T cell interactions in shaping the immune response in the metastatic microenvironment.

**Fig. 2:**
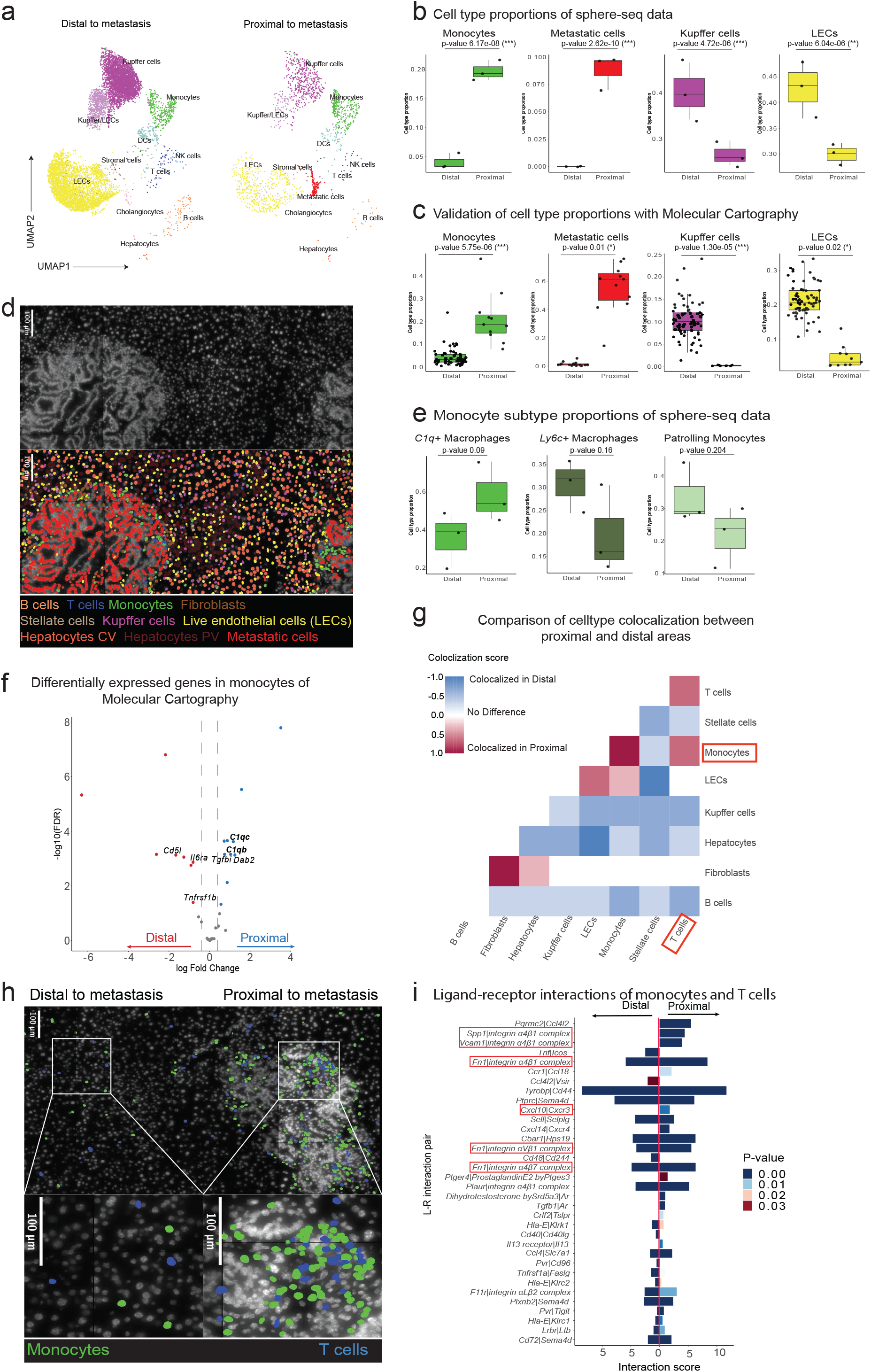
Sphere-sequencing of metastases-bearing mouse liver reveals cell types and interactions specific to the metastatic-niche. **a**, Uniform manifold approximation and projection (UMAP) visualization of integrated samples of mouse metastatic liver (n=3 samples, which only includes samples with a high number of metastatic cells). Cells are clustered, annotated and coloured by their cell type. Cells are separated based on the amount of metastatic cells in their sphere of origin (distal = spheres without metastatic cells; proximal = spheres with metastatic cells). **b**, Boxplots of selected cell type proportions of grouped sphere-seq data in distal and proximal regions (n=3 samples). From left to right; monocytes, metastatic cells, Kupffer cells (KCs) and liver endothelial cells (LECs). Black dots represent individual mice. **c**, Validation of cell type proportions with Molecular Cartography. Displayed like in b, but the y-axis shows cell type proportion per spatial area and black dots represent spatial areas (n= distal: 71 areas, proximal: 11 across 2 samples). **d**, Representative Molecular Cartography image; upper image, DAPI stain (white), lower image, DAPI stain overlayed with cell type annotations. **e**, Boxplots of cell type proportions from sphere-seq showing monocyte subtypes (*C1q+* macrophages, *Ly6c+* macrophages and patrolling monocytes) (n=3 samples). **f**, Differentially expressed genes (DEGs) of monocytes between distal and proximal regions from Molecular Cartography highlighted in a volcano plot (n= distal: 71 areas, proximal: 11 across 2 samples). Colored dots represent significantly enriched genes; blue, enriched in proximal; red, enriched in distal. **g**, Cell colocalization map built from Molecular Cartography data comparing frequency of colocalization in distal and proximal areas (n = 2 samples). For calculation of colocalization score see methods. **h**, Molecular Cartography images with monocytes highlighted in green and T cells in blue comparing distal and proximal metastatic sites. **i**, Predicted ligand-receptor (L-R) interactions between monocytes and T cells in distal or proximal areas based on sphere-seq data (n=3 samples). Interaction scores were calculated by CellPhoneDB, which uses a permutation test to generate p-values indicating significantly enriched L-R interactions. Interactions referenced in the main text are highlighted with red squares. For **b**, **c**, **e** and **f** a negative binomial generalized log-linear model was used for statistical testing and p-values (benjamini-hochberg adjusted) of <0.05 were considered significant (***<0.001, **<0.005, *<0.05).

To identify potential drivers of these interactions we performed L-R interaction analysis^13^ which revealed a number of spatially restricted interactions (Fig. 2i). Notably, this included interactions of Secreted Phosphoprotein 1 (*Spp1*), Fibronectin-1 (*Fn1*) and *Vcam1* with α4β1 integrin complexes on T cells (Fig. 2i, Supplementary Fig. 6b). While VCAM-1 α4β1 interactions are mostly noted as adhesive interactions, SPP1 and FN1 are associated with a poor prognosis in various cancer types^16–18^. SPP1 was previously found to suppress T cell responses in the tumor microenvironment^19^, while FN1 reportedly correlates with infiltration of anti-inflammatory macrophages^18^. Interactions of both of these monocyte-derived ligands with integrins on other immune cells were found to be enriched in colorectal cancer and might be involved in promoting tumorigenesis^20^. In addition, we found a proximal enrichment of *Cxcl10|Cxcr3*, an interaction which is well-known for mediating effector T cell recruitment^21^ and has been found to be enriched between *C1q+* macrophages and other immune cell subsets in colorectal cancer^20^.

### Sphere-sequencing can be adapted to other tissue types and species

Sphere-seq can be easily adapted to other tissue types and species with minor adjustments to single cell dissociation. We have created proof-of-principle datasets from the mouse spleen and further showed that the method is compatible with clinical samples by applying it to Crohn’s disease biopsies (Supplementary Fig. 7a-d,). For clinical samples, we introduced size-selection based on sequential filtering and manual picking of spheres to demonstrate the feasibility of the protocol in the absence of an expensive biosorter apparatus (Supplementary Fig. 7e).

## Discussion

Like any spatial transcriptomics methodology, sphere-seq has inherent strengths and limitations. Direct comparison to Visium^22^, one of the most widely used spatial transcriptomics methods, demonstrates the strength of sphere-seq. Visium’s *in situ* permeabilization protocols need to be optimized for different tissue compositions, resulting in compromise solutions and uneven data quality in heterogeneous tissues such as metastasis-bearing organs, while sphere-seq demonstrates far more consistent results (Supplementary Fig. 8a-c). The greatest advantage of sphere-seq compared to slice-based spatial transcriptomics is single cell resolution. In contrast, Visium data requires deconvolution of spots into single cells, which revealed a clear bias towards hepatocytes in our datasets (Supplementary Fig. 8d). This phenomenon has been previously described^23^ and is attributed to the RNA content of hepatocytes superseding that of smaller cells like LECs.

In our dataset, Visium failed to reliably detect the LEC and KC zonated genes *Plpp1, Galnt15*, and *Vcam1*, therefore, we assessed their expression in publicly available Visium datasets from healthy wild type and NAFLD (non-alcoholic fatty liver disease) mice^10^. This confirmed zonated gene expression, with *Vcam1* expression being more heavily expressed and zonated in NAFLD compared to healthy livers, in line with its involvement in inflammatory processes^11^ (Supplementary Fig. 8e,f). However, due to the lack of single cell resolution of Visium data, it cannot be directly determined whether zonated gene expression is caused by KCs or LECs. In fact, prior knowledge of the importance of *Vcam1* in LECs would likely lead to neglecting its zonation in KCs. In sum, comparison with Visium demonstrates sphere-seq’s superior performance in uncovering zonated gene-expression patterns in subdominant cell types. As demonstrated here, sphere-seq enables unsupervised spatial hypothesis generation for entire tissues. This makes sphere-seq an ideal method for exploratory research, which can then be validated by image-based spatial transcriptomics or proteomics methods that offer subcellular spatial resolution but require prior knowledge of gene expression for panel assembly.

Another advantage of sphere-seq is its compatibility with existing downstream protocols used for scRNA-seq, such as the vast existing catalog of computational methods. Furthermore, sphere-seq has the potential to profile other modalities, such as chromatin accessibility or proteins, solely by adapting the scRNA-seq protocol to ATAC-seq^24,25^ or CITE-seq^26^ approaches, respectively. So far, sphere-seq has been exclusively applied to fresh tissues, however, the protocol could be modified for frozen tissues. Sphere-seq could theoretically be adapted to single nuclei RNA sequencing by modifying lysis protocols and exchanging lipid-with cholesterol-tagging barcodes^5^. Like any methodology that uses spatial reconstruction, sphere-seq is limited by prior knowledge of landmark gene expression patterns^27^ to reconstruct the position of spheres within tissues. This could be resolved by combining sphere-seq with array-based spatial transcriptomics methods to identify spatial landmark genes and group spheres according to their spatial niches. Unlike sliced-based spatial transcriptomics methods, sphere-seq is not limited to individual two-dimensional regions of interest, and is therefore more representative of the sample tissue as a whole. However, spheres are generated in a random process and rare niches might not be reliably captured. This could be amended by introducing fluorescently-labeled antibodies or fluorescent reporter genes to indicate rare niches of interest and select for signal-positive spheres with the large sample sorter.

In sum, we have shown that sphere-seq is a powerful new tool to investigate differences in cellular composition and gene expression between distinct tissue niches. To the best of our knowledge, we here describe zonated gene expression in KCs for the first time. Our new method furthermore empowered us to identify L-R interactions which are not only significantly enriched in the dataset but are also demonstrated to co-occur spatially. Finally, we showed that sphere-seq can be applied to other tissues and species. Altogether, we show that sphere-seq is a valuable addition to currently available technologies in the field of spatial transcriptomics.

## Acknowledgments

We thank Owen Sansom (Beatson Institute for Cancer Research, Glasgow, UK) for the AKPS organoids and Salvatore Piscuoglio (Department of Biomedicine, University of Basel, Switzerland) for the human colon cancer organoids. We also thank Michael Scharl and Barbara Maria Szczerba (Gastroenterology and Hepatology, University Hospital Zürich, Switzerland) for collection of Crohn’s disease biopsies and we thank the patients and their families for participating in this study. For help and support with the large fragment biosorter we thank Sven Panke and Steven Schmitt (Department of Biosystems Science and Engineering, ETH, Basel, Switzerland) and Erich Brunner (Department of Molecular Life Sciences, University of Zurich, Switzerland). We thank Laura De Vargas Roditi (Department of Biosystems Science and Engineering, ETH, Basel, Switzerland) for interesting discussions for analysis pipeline development. And finally, we thank Zev J. Gartner (UCSF, San Francisco, USA) for providing us with lipid anchor and co-anchor (MULTI-seq) and Christopher S. McGinnis (UCSF, San Francisco, USA) for technical advice. We used BioRender for creating schematic drawings. This work was funded by the Swiss National Science Foundation (PCEFP3_181249 to A.E.M.) and the Leona M. and Harry B. Helmsley Charitable Trust (Gut Cell Atlas grant to A.E.M.).

## Author contributions

K.H. and C.B. performed the experiments. K.H., K.B. and I.E.A. performed the data analysis. A.M. supervised the study. K.H. and A.M. wrote the paper. X.F. contributed to editing and writing the paper. All authors discussed the results and commented on the manuscript.

## Material and Methods

### Mouse model of liver metastasis

VilCreERT2;APCfl/fl;Tp53fl/fl;KrasG12D/wt (AKP) organoids, obtained from Owen Sansom from Beatson Institute for Cancer Research in Glasgow, were modified with an additional knockout in Smad4, resulting in VilCreERT2;APCfl/fl;Tp53fl/fl;KrasG12D/wt;Smad4KO (AKPS). Organoids were cultured in Matrigel (Corning) as described previously^28^. The culture medium (Advanced DMEM/F12, Life Technologies™) was supplemented with 10 mM HEPES (Life Technologies™), 2 mM L-Glutamine (Life Technologies™), 100 mg/mL Penicillin/streptomycin, 1x B27 supplement (Life Technologies™), 1 x N2 supplement (Life Technologies™) and 1mM N-acetylcysteine (Sigma-Aldrich). For splenic injections, AKPS organoids (4 domes per mouse) were repeatedly washed in ice cold PBS to remove all Matrigel, then mechanically dissociated into small fragments, resuspended in 50 uL PBS and loaded in an insulin syringe (BD, MicroFine, 0.3 ml, 30 G). C57BL/6 mice were administered Carprofen (5 mg/kg) subcutaneously 30 min before surgery, then anesthetized with isoflurane gas and kept warm on a 37 °C thermal pad. After shaving and disinfection with betadine, a mixture of 0.5% lidocaine (5 mg/ml) and 0.25% bupivacaine (2.5 mg/ml) was subcutaneously injected along the planned incision line. The spleen was exposed and the organoids were injected under the splenic capsule. After 10 minutes, the spleen was resected by ligation. The wound was washed with sterile PBS, the peritoneal wall was closed with an absorbable polyglactin suture (Vicryl 4-0 or 5-0 coated) and the skin with wound clips. Buprenorphine (0.1 mg/kg) was injected subcutaneously during the wake-up phase. The animals were monitored until awake and in the days following surgery. The experiment was terminated 14 days after intrasplenic injection. All experimental procedures were performed in accordance with Swiss Guidelines and were approved by the Cantonal Veterinary Office Basel-City.

### Colon cancer organoids mixing species experiment

GFP+ mouse and GFP-human colon cancer organoids were mixed in a ratio of 1:1 and used for sphere-seq. Based on the GFP signal 144 wells were sorted with GFP+ mouse and 144 wells with GFP-human organoids.

#### Cultivating human colon cancer organoids

Human colon cancer organoids were obtained from the Visceral Surgery Research Laboratory lead by Salvatore Piscuoglio at University of Basel and cultured in Matrigel (Corning) as described previously^28^. The culture medium (Advanced DMEM/F12, Life Technologies™) was supplemented with 10 mM HEPES (Life Technologies™), 2 mM Glutamax (Gibco), 1x B27 supplement (Life Technologies™), 10 mM Nicotinamide (Sigma Aldrich), 1.25 mM N-acetyl-l-cysteine (Sigma Aldrich), 500 nM A83-01 (Stemcell #100-0245), 50 ng/ml Human epidermal growth factor (hEGF) (Stemcell #78006.1), 100 ng/ml Recombinant human Noggin (Stemcell #78060), 100 ng/ml human R-spondin (LuBioScience #120-38-20), 10 nM Prostaglandin E2 (PGE2) (Stemcell #72634), 10 μM SB202190 (Stemcell #72634), 10 μM Y-27632 dihydrochloride (Rock inhibitor) (Stemcell #72304) and 10 nM Gastrin (Sigma Aldrich, #G9145).

##### Cultivating mouse colon cancer organoids

VilCreERT2;APCfl/fl;Tp53fl/fl;KrasG12D/wt (AKP) organoids obtained from Owen Sansom were labeled with the plasmid pMSCV-loxp-dsRed-loxp-eGFP-Ruo-WPRE^29^ and cultured as described in ‘Mouse model liver metastasis’ section with the addition of 100 μg/ml murine recombinant Noggin to the culture medium (LuBioScience, #250-38-250).

#### Tissue collection

##### Liver

Mice were euthanized and the liver was perfused with PBS (pH 7.4, Gibco) at a flow rate of 2-3 mL/min over an insertion of the cannula in the inferior vena cava. After the liver was fully perfused the gallbladder was removed and the liver was collected in ice cold PBS. For Molecular Cartography and Visium a piece of the liver was embedded in O.C.T.™ compound (Tissue-Tek) and snapped frozen in a metal beaker filled with isopentane (Sigma Aldrich) in liquid nitrogen. Snap-frozen tissues were stored at −80 °C.

##### Spleen

Mice were euthanized, the spleen was removed and collected in ice cold PBS.

##### Crohn’s Biopsy

Tissue was surgically resected and collected in MACS tissue storage solution (Miltenyi Biotec #130-100-008) on ice for around 4-6 hours before further processing.

##### Colon cancer organoids

Matrigel of domes was dissolved with ice cold PBS and organoids were collected in ice cold PBS after shaking on ice for 30 min to remove all residual matrigel.

#### Sphere-sequencing workflow

##### Partial dissociation into spheres

Tissues were partially dissociated into spheres using the mechanical force of a scissor. Spheres were then first filtered with at 400 μm strainer (PluriSelect #43-50400-03) to remove larger spheres that would clog the biosorter. Then spheres were filtered a second time using a 40 μm strainer (PluriSelect #43-50040-51) to remove single cells. Thereby, after solution was filtered through the 40 μm strainer, the strainer was turned and larger than 40 μm spheres were washed off the strainer and collected in a 50 ml falcon tube in PBS. Liver spheres were collected and sorted in low glucose DMEM (1g/L D-Glucose/L-Glutamine, Pyruvate, Gibco #31885-023). The same filtering approach was used for Crohn’s disease biopsies with a lower size strainer of 200 μm and a higher size strainer of 500 μm resulting in a suspension of spheres between 200-500 μm. Solution (Advanced DMEM/F12, Life Technologies™) with spheres was then put into a petri dish and spheres were manually picked under a stereomicroscope (Leica) using a P200 pipette and tips.

##### Sphere sorting using a large fragment biosorter (Copas)

Spheres were sorted into 96 well plates (non-binding, v-shaped, Greiner bio-one #651901) filled with 30 μl dissociation medium, one sphere per well using a large fragment biosorter (Copas) with a 1000 μm large nozzle. The gates were set to allow sorting of spheres within specific size ranges between 200-400 μm. The size was defined with a linear model by acquiring time of fight (TOF) of standard sized beads (Megabead NIST Traceable Particles (Polysciences): 60 μm (#64200-15), 125 μm (#64225-15) and 175 μm(#64235-15)). TOF on the x-axis and extinction of the y-axis measures were then predicted for sizes between 200-400 μm from the linear model to gate and sorted accordingly. For the organoids species mixing experiment GFP+ mouse organoids were sorted using the 488 nm blue laser at a power of 50. Gates for positive ones were drawn against a GFP-organoid control with similar organoid size. Solution with spheres was diluted to around 10 events per second for proper sorting with pure/no doublets sorting mode. Other settings were the following: Power 50 mW; gain 1.0; PMT Volts green 600, red 750; drop width 7 mS; sort delay 23 mS; sample cup pressure 0.42 psi; diverter pressure 2.5 psi; sheath flow rate 57 %. Sheath flow solution was PBS (pH 7.4, Thermo Fisher).

##### Single cell dissociation

Spheres were sorted directly into dissociation buffers. Spleen dissociation buffer was PBS (pH 7.4, Gibco). Spheres were sorted into 30 μl of PBS and dissociated by applying mechanical force pipetting up and down around 50 times using a P20 multichannel pipette. Liver dissociation buffer was a mix of low glucose DMEM (1g/L D-Glucose/L-Glutamine, Pyruvate, Gibco #31885-023), 15mM HEPES (Gibco) and 32 μg/ml liberase™ (Sigma Aldrich #05401119001) and 1x TripLE (diluted from 10x TripLE) (Gibco). Plates with dissociation buffers of 30 μl and sorted spheres were incubated at 37 °C shaking at 300 rpm for 20 min. The Crohn’s biopsies spheres were sorted into 100 μl epithelial dissociation medium (HBSS(-Ca2+-Mg2+) (Sigma Aldrich), 10mM HEPES (Gibco) and 5mM EDTA (Lonza)) and incubated at 37 °C and 300 rpm for two times 15 min with vortexing of plates in between for 30 s. After incubation plates were spun at 400 xg for 10 min at 4 °C and the supernatant was carefully removed after. Then 100 μl of digestion medium (HBSS(+Ca2+, +Mg2+) (Sigma Aldrich), 0.5 mg/ml DNAseI (Roche Diagnostics #10104159001) and 0.5 Collagenase from Clostridium histolyticum (Sigma Aldrich #C5138)) were added and incubated at 37 °C and 300 rpm for 30 min. After incubation enzyme activity was inactivated by adding 50 μl of 20 mM EDTA/PBS (Lonza, pH 7.4 Gibco) shaking at 300 rpm 37 °C incubating for 5 min. Then plates were spun at 4 °C 400 rpm for 10 min and supernatant except around 30 μl was removed after. Colon cancer organoids were sorted into 30 μl of 1x TripLE (Gibco) and incubated at 37 °C for 12 min at 300 rpm.

##### Labeling of cells with sphere specific barcodes

For labeling of spheres the MULTI-seq lipid hashing method was used^5^. Lipid anchor and co-anchor were obtained from Zev J. Gartner laboratory at University of California San Francisco. We designed a set of 288 barcodes with a minimum hamming distance of 3 using the Bioconductor package DNAbarcodes^30^ (Version 1.20.0), incorporated primer binding sequences from the MULTI-seq method^5^ and ordered them with a purity of standard desaulting. Anchor and barcodes were mixed beforehand in a volume of 20 μl in PBS in a concentration of 50 nM for spleen, liver and organoids and 100 nM for Crohn’s biopsies in a ratio of 1:1. Co-Anchor were also diluted beforehand in PBS in the same concentrations.. After dissociation of spheres 20 μl of Anchor:Barcode mix were added to the wells and mixed for 30 s 700 rpm on a thermomixer at 20 °C. Then the cells were incubated on ice for 5 min. After incubation the diluted co-anchor is added and again mixed for 30 s 700 rpm at 20 °C followed by an incubation on ice for 5 min. To quench binding of lipids to the cells 100 μl of 10 % BSA (Sigma Alrich) in PBS were added, mixed for 30 s 700 rpm and incubated on ice for 5 min. After incubation cells from all wells were pooled into FACS tubes (Falcon) and spun at 400 xg for 10 min for Crohn’s biopsy and spleen, 300 xg for 10 min for liver and 300 xg for 5 min for colon cancer organoids. Cells were then washed at least 2-3 times with 1 % BSA in PBS. For the last wash cells were transferred into a 1.5 ml DNA lobind tube (1.5 ml, Eppendor) and spun a last time to then resuspend cells in only around 50-100 μl for counting and quality check using a hemocytometer and trypan blue solution (0.4 %, Thermo Fisher). Samples were processed further for scRNAseq if there were at least 10,000 cells and the viability of cells was at least 70 %.

#### Single cell RNAseq library preparation using BD Rhapsody and MULTI-seq

Whole Transcriptome Analysis (WTA) on MULTI-seq labeled cells was performed using the BD Rhapsody Single-Cell Analysis System (BD Biosciences). In total 9 samples were processed of mice that were injected with colon cancer organoids, one sample of healthy mouse liver, one sample of mixed colon cancer organoids and 2 samples of Crohn’s disease biopsies. For each sample a BD Rhapsody cartridge was loaded with approximately 10,000 cells. Single cell capture and cDNA synthesis using the Single Cell Capture and cDNA Synthesis kit (#633731, #633733, #633773) were done following the manufacturer’s protocol (BD Biosciences). Libraries were then prepared using the BD Rhapsody WTA Amplification Kit (#633801) following instructions of the mRNA WTA Library Preparation Protocol (BD Biosciences). For the MULTI-seq library first the BD protocol of Sample Tag Library Preparation was followed until purification of Sample Tag PCR1 product with the difference of adding the MULTI-seq primer (sequence according to McGinnis and Patterson et al.^5^) in a concentration of 10 μM instead of Sample Tag PCR1 Primer for Sample Tag PCR1 reaction. After purifying the Sample Tag PCR1 product indexing PCR was done following instructions of the MULTI-seq protocol where small RNA TrueSeq indexing primers (Illumina #15004197) were used for i7 and the Forward Primer from the BD WTA kit was used for i5.

#### Single cell RNAseq library preparation using Chromium 10X and MULTI-seq

Libraries were generated following manufacturer’s instructions from Chromium Next GEM Single Cell V(D)J Reagent Kits v1.1 protocol. In short: cells were resuspended in 0.04 % BSA and mixed with Master mix containing reagents for reverse transcription. Cell suspension was then loaded in GemCode Single-cell Instrument (10X Genomics) together with GemCode Single-Cell 5’ Gel Beads. Cells and beads were fused to generate single-cell Gel Bead-in-Emulsions (GEMs). Within GEMs cells were lysed and RNA was reverse transcribed. After GEMs were broken and cDNA was cleaned up, using DynaBeads MyOne Silane Beads (Thermo Fisher #37002D) and SPRIselect beads (Bekman Coulter #B23318), cDNA was amplified and cleaned up using SPRIselect beads. Then amplified cDNA was enzymatically fragmented and indexed sequencing libraries were generated by the following steps: end repair, A-tailing, adapter ligation, post-ligation SPRIselect cleanup and sample index PCR. For MULTI-seq library preparation instructions were followed from McGinnis and Patterson et al.^5^. In short: The MULTI-seq primer (according to MULTI-seq protocol instructions) was added in a concentration of 10 μm to the cDNA amplification mix. After cDNA amplification during SPRIselect cleanup the non-bound fraction (containing small cDNA fragments) was saved and cleaned up with SPRIselect beads in a higher ratio to enrich for small MULTIseq barcodes. These products were then used for index PCR using the SI-PCR Primer from the 10X kit for the i5 and one of the small RNA TrueSeq index primers for the i7 (Illumina #15004197).

#### Visium library preparation

Two mouse liver samples with visible micro-metastasis were processed. A 10X Visium Spatial Gene expression slide was put into the cryostat (Leica CM3050S) to calibrate its temperature to −20 °C. Then 10 μm sections of metastatic mouse liver samples were cut and placed within the capture area. The capture slide was then stored in a slide container at −80 °C until the next day for further processing. cDNA libraries were generated following the manufacturer’s instruction. In short: Tissues were fixed with methanol and hematoxylin and eosin (H&E) staining was done to check tissue quality and morphology. Then tissue lysis, reverse transcription, second strand synthesis and cDNA denaturation were performed on the slides. Permeabilization time of 10 min was assessed beforehand with the Tissue Optimisation Protocol. Reactions were transferred into PCR tubes and qPCR was done to measure cDNA concentration. cDNA was then amplified by PCR using cycle numbers defined by qPCR. Final library preparation steps (End repair, A-tailing, adapter ligation and sample index PCR) were done to generate indexed sequencing libraries.

#### Quality assessment of libraries and sequencing

Quality and quantity of all libraries were assessed using the dsDNA high-sensitivity (HS) kit (Life Technologies #Q32854) on a Qubit 4 fluorometer (Thermo Fisher) and the high sensitivity D1000 reagents and tapes (Agilent #5067-5585, #5067-5584) or high sensitivity D5000 reagents and tapes (Agilent #5067-5593, #5067-5592) on a TapeStation 4200 system (Agilent Technologies). Paired-cell sequencing was performed for all libraries (WTA BD libraries: read 1: 60bp, index read: 8bp, read2: 62-100bp; WTA 10X libraries: read 1:26bp, index read: 8bp, read2: 88-96bp; MULTI-seq libraries: read 1: 26bp (10X), 60bp (BD Rhapsody), index read: 6 bp, read2: 62-100bp)) on a NovaSeq 6000 system (Illumina) using NovaSeq SP Reagent Kits (100 cycles) v1.5 and S4 Reagent kits (200 cycles) v1.5 with XP workflow. WTA libraries of BD Rhapsody and 10X were sequenced 50,000 reads/cell, MULTI-seq libraries of both 5,000 reads/cell and Visium libraries 50,000 reads/spot.

#### Single cell RNAseq and Visium data preprocessing

##### Demultiplexing

BCL files were demultiplexed using Bcl2fastq v2.20.0.422 from Illumina to convert them to FASTQ files.

##### BD Rhapsody data was preprocessed using zUMIs

FASTQ files from WTA libraries of BD Rhapsody data were processed using the zUMIs^31^ (v2.9.4) platform to convert reads to count matrices per cell. For gene alignment STAR^32^ (v2.5.2b) was used with the following genecodes: for human Crohn’s disease biopsy samples GRCh38 v2020-A; for mouse liver samples GRCm38 vM25 fused with GFP 3’ UTR sequence and for the colon cancer organoids mixing species experiment a fused index of GRCm38 v2020-A, GRCh38 v2020-A and GFP 3’ UTR sequence. Three liver samples (S1-3) were sequenced twice to acquire deeper sequencing, fastq files for read1 and read 2 were merged and the merged files were used as zUMI input.

##### 10X data was pre-processed using Cell Ranger

Mouse spleen data single cell count matrices were generated using Cell Ranger (v5.0.0) (10X Genomics) with GRCm38 v2020-A genecode.

##### Visium data was pre-processed using Space Ranger

Mouse liver Visium data was pre-processed using Space Ranger (v1.2.0) (10X Genomics) with GRCm38 v2020-A genecode.

#### Sphere-sequencing downstream analysis

Downstream analysis of UMI count matrices was done in R version 4.1.0 and most analysis was done using the following packages: Seurat^33^ (v4.0.3), scran^34^ (v1.22.1) and SingleCellExperiment^35^ (v1.16.0). Dplyr^36^ (v1.0.7) and tidyverse^37^ (v1.3.1) were used for data wrangling. Plotting was mostly done with ggplot2^38^ (v3.3.5).

##### Conversion of genecode numbers from zUMI outputs to gene names

Genecode numbers were converted using the biomaRt^39,40^ Bioconductor package (v2.50.3) with musmusculus_gene_ensemble version 95 for mouse data and hsapiens_gene_ensembl version 95 for human data. For the organoid species mixing experiment both genecodes were used. GFP gene name was included.

##### Sphere barcode classification and integration with whole transcriptome analysis (WTA)

For allocation of MULTI-seq barcodes to single cells the deMULTIplex^5^ (v1.0.2) workflow on the github repository https://github.com/chris-mcginnis-ucsf/MULTI-seq was followed. Briefly, a sample barcode UMI matrix per cell was generated and then cells were assigned to specific barcodes following the classification workflow. Cell barcodes of classified cells were then matched with cell barcodes of WTA.

#### Mouse liver specific sphere-seq downstream analysis

##### Normalization and batch effect correction

All 10 liver samples were merged after generation of Seurat objects and UMI (unique molecular identifier) counts underwent SCT normalization (Seurat function, ‘SCTransform’) which normalizes and scales data and finds variable features. Then the merged Seurat object was transformed into a SingleCellExperiment object and batch effect correction was done using MNN (mutual nearest neighbors) correction within the batchelor^41^ Bioconductor package (v1.10.0).

##### Quality control and clustering

After batch effect correction low quality cells (lower than 200 features and higher than 20% of reads mapped to mitochondrial genes) and doublets (higher than 7,500 features) were removed and 10 MNN corrected principal components (PCs) were used for clustering of cells in Uniform Matrix Approximation and Projection (UMAP) two-dimensional space.

##### Cell type annotation

Initial clustering with 10 MNN corrected PCs and a resolution of 1 was first broadly annotated using cell type markers from the liver cell atlas^10^. Then each broadly annotated cell cluster was further investigated for subtypes using the Seurat subclustering function ‘FindSubCluster’. Subclusters were then annotated with help of the liver cell atlas and investigation of differentially expressed genes (DGE) using the Seurat function ‘FindAllMarkers’ with a non-parametric Wilcoxon Rank Sum test with default parameters (min.pct 0.25 and logfc.threshold 0.25).

##### Integration of sphere size from biosorter data

A standard curve was generated with TOF measurements of standard sized beads. TOF measurements of sorted spheres were then fitted into the linear model to calculate the size in diameter.

##### Cells per sphere cutoff

For further analysis only spheres with at least 5 cells were used.

##### Lobule layer classification

For each sphere a zonation coordinate (ZC) based on zonated landmark genes in LECs was calculated as previously described^42^. Briefly, pseudo bulks from LECs of each sphere were generated using central and portal landmark genes from Halpern *et al*.^2^ Genes were normalized by dividing its expression by the maximum level of expression across spheres to ensure equal contribution of all genes. Then the sum of central and portal landmark genes (cLM and pLM) was calculated for each sphere and used to calculate ZCs by the following calculation: ZC = pLM/(pLM+cLM). At the end ZCs were rescaled so that 0 is the most central and 1 is the most portal coordinate: ZC = (ZC - min(ZC))/(max(ZC) - min(ZC)). Spheres without LECs were removed from further analysis at this point. ZCs of spheres were then grouped into lobule layers L1-L10 (L1: ZC <0.1,L2: ZC <0.2,L3: ZC <0.3,…,L10: 0.9< ZC ≤1.0) There were not many spheres from the most central and most portal areas so they were grouped into L1-L3 and L8-L10. To control the zonation algorithm in our data, landmark genes in hepatocytes were analyzed from spheres grouped into central (L1-L5) and portal (L6-L10) veins.

##### Split of dataset for different analysis

After pre-processing analysis data was split into three datasets. The healthy liver sample (n=1), the liver samples of mice that were injected with colon cancer organoids (n=9) for analyzing liver zonation during metastasis formation and liver samples with a high amount of metastatic cells (at least 20 within all cells) (n=3, samples S3, S6 and S7) for analysis of different metastatic niches (Supplementary Fig. 6a).

##### Metastatic distance classification

Spheres with metastatic cells were grouped into ‘proximal’ and spheres without metastatic cells into ‘distal’ categories

##### Analysis of zonation specific genes

Differential gene expression analysis was performed using edgeR^43–45^ (v3.36.0). For this single cell counts were summed across spheres with at least 5 cells of a cell type of interest to derive a single expression vector per sphere. After removing lowly expressed genes using the ‘filterByExpr’ function in edgeR, a negative binomial generalized log-linear model was fitted to the remaining genes with the lobule layer as an ordered factor covariate (L1-L3<L4<L5<L6<L7<L8-L10). The linear coefficients were then used for fitting and the sample names were used as a blocking factor to account for batch effects. The ‘glmQLFTest’ function was used to identify genes with coefficients for the linearly encoded factor significantly different from 0 at a Benjamini-Hochberg adjusted p-value of 0.05. This analysis was done for LECs and KCs. In LECs only genes that were not in the landmark gene panel to calculate ZCs were further investigated.

##### Differential gene expression (DGE) analysis between two groups

DGE analysis was done like described above with two differences. First, single cell counts were summed across spheres with at least 2 cells of a cell type of interest, and second instead of using lobule layers as an ordered factor covariate, two groups were used as factor covariates.

##### Analysis of differences in cell type abundance between two groups

To identify changes in cell-type-specific abundance between two groups, normalized log counts of cluster abundance were computed using the ‘cpm’ function in edgeR^43–45^ (v3.36.0) accounting for total number of cells per sample^46^. After specifying a design matrix with group labels (veins ‘central’ and ‘portal’ or distances to metastasis ‘proximal’ and ‘distal’) as covariates and sample names as blocking factors, the dispersion parameter of the negative binomial model was estimated using the ‘estimateDisp’ function in edgeR with trend=‘none’. a negative binomial generalized log-linear model was fitted with ‘glmQLFit’ function (robust=TRUE, abundance.trend= FALSE) for each cell type. The ‘glmQLFTest’ function was then used to identify cell types with coefficients significantly different from 0 at a Benjamini-Hochberg adjusted p-value of 0.05.

##### Ligand-receptor (L-R) interaction between different groups using CellPhoneDB

L-R interaction analysis was done using the Python Package CellPhoneDB^13^ (v4.0.0) following instructions on the GitHub repository https://github.com/ventolab/CellphoneDB. In short, gene expression of annotated clusters was used as input to match with known L-R interaction pairs from the CellPhoneDB public repository using default parameters. The average ligand and receptor expression between two cell types were represented by the mean values which were calculated using the percentage of cells within a cluster expressing the ligand or receptor and their gene expression mean. A null distribution of means for randomly permuted annotated cluster labels was then used to determine p-values. Analysis was done for separate groups of veins (central and portal) and distances to metastasis (proximal and distal). Differences in L-R interaction scores between two groups were then visualized in a barplot with L-R pairs ordered decreasing by the difference in interaction scores between two groups.

#### Colon cancer organoids mixing species specific downstream analysis

##### Integration of GFP fluorescent signal from biosorter data

Sphere size was calculated with help of a standard curve from standard sized beads. The GFP signal was then normalized by dividing it with the sphere-size to account for autofluorescence that is higher in larger spheres.

##### Quality control, normalization, clustering and annotation

Low quality cells were removed with lower than 200 features and larger than 30% of reads mapped to mitochondrial genes. UMI counts were then normalized and scaled using the ‘SCTransform’ Seurat function and 10 PCs were used for clustering in UMAP space. Cell clusters were then annotated as human or mouse depending on the species of genes being expressed. Based on the knowledge of sorting, cells could also be annotated by their species-well.

##### DecontX to remove cell free RNA

During data exploration of human and mouse UMI reads per cell we found a lot of cell free RNA in sphere-seq data even after removal of low quality cells. This was probably due to the low quality of colon cancer organoids (~20-30% dead cells before single cell capture). Therefore decontamination of data was done using the decontX function (default parameters) within the celda^47^ R package (v1.12.0).

##### Analyzing the fraction of correctly and wrongly assigned cells

There are two annotations, the cell species annotation and the well annotation that is established by sorting of a human or mouse organoid. By matching these two pieces of information the proportion of wrongly and correctly assigned cells for each sphere could be analyzed, wrong if there were mouse cells in human wells and human cells in mouse wells, correct if the species was matching. Three spheres were found to be 100% made out of wrongly assigned cells, and were therefore allocated to the opposite species well because these are most probably due to a fluorescence sorting error.

#### Mouse spleen specific downstream analysis

##### Quality control, normalization, clustering and annotation

Two samples were merged after sphere barcode integration; no batch effect correction was needed. Low quality cells were removed, with lower than 200 features and larger than 10% of reads mapped to mitochondrial genes, and doublets were removed with larger than 6000 features. UMI counts were then normalized and scaled using the ‘SCTransform’ Seurat function and 10 PCs were used for clustering in UMAP space. Cell clusters were annotated using cell type marker genes from Medaglia *et al*.^48^. Only spheres with at least 5 cells were considered for plotting.

#### Crohn’s disease biopsy specific downstream analysis

##### Quality control, normalization, clustering and annotation

Two samples were merged without the need of batch effect correction. After low quality cells and doublets (cells with lower than 200 features, larger than 25% of mitochondrial reads, larger than 5000 features) were removed, UMI counts were normalized and scaled (‘SCTransform’) and 10 PCs were used for clustering. Cells were annotated using marker genes from Martin *et al*^49^. At least 5 cells per sphere were required for plotting.

#### Visium data downstream analysis

##### Normalization, clustering and spatial area annotation

Samples were processed separately, normalization and scaling was done using ‘SCTransform’ Seurat function and clustering was done using 10 PCs in UMAP space. Clusters were then annotated into the following areas: portal and central veins (also considered as ‘distal’ to metastatic sites), and metastasis (considered as ‘proximal’ to metastatic sites) based on landmark genes.

##### Comparison of number of gene features

Mean values of the number of gene features between proximal and distal metastatic areas of Visium data were compared with mean gene feature values of both areas from sphere-seq data. The same two metastatic liver samples were used.

##### Batch effect correction

The two Visium samples were merged and batch effect correction was done using MNN (mutual nearest neighbors) correction within the batchelor^41^ Bioconductor package (v1.10.0).

##### Deconvolution

For deconvolution of spots, annotated sphere-seq data was used as a reference. The top 20 genes per cell type cluster were selected that were also expressed in Visium datasets. These genes were used for deconvolution using the SCDC^50^ package (v0.0.0.9000). Spots were allocated to a specific cell type if at least 75% of genes could be assigned to one cell type. Spots with less than 75% are annotated as mixed.

##### Public Visium data analysis

Public Visium data from Guilliams *et al*.^10^ was used from wild-type mouse and NAFLD (non-alcoholic fatty liver disease) mouse models. Spots annotated for different liver zones (central, mid, periportal and portal) were grouped and tested for gene expression of newly found zonation specific genes.

#### Highly multiplexed FISH (Molecular Cartography™)

##### Sample preparation, Probe design, Imaging and Pre-processing

These steps were done as previously described^10^. In brief, liver samples (4 mouse liver metastasis samples of which two samples had visible micrometastasis) were frozen and sectioned to 10 μm slices as described for Visium; sections were placed within capture areas on Resolve BioScience slides. After, slides were sent to Resolve BioSciences on dry ice, where they were processed further. Samples were fixed and underwent 100-plex combinatorial single molecule fluorescence in-situ hybridization. During multiple cycles of color development, imaging and decolorization a unique combinatorial code for each target gene was generated. 100 genes were chosen from sphere-seq analysis (20 genes were chosen to define cell types and 80 genes were hits to validate) and their probes were designed using Resolve’s proprietary design algorithm that makes sure that probes are specific with little off-target binding. Fluorescent signals were imaged on a Zeiss Celldiscoverer 7 microscope with a final magnification of 25x. Each region underwent 9 imaging rounds and 16 z-stacks were acquired. Java and C++ scripts were then used for spot segmentation and images were pre-processed to remove background fluorescence. Raw data images from different imaging rounds were aligned during which images had to be corrected using an iterative closest point cloud algorithm. Then a profile for each pixel was created using the information of 16 values (16 images from two color channels in 8 imaging rounds).

##### Downstream analysis

Image analysis was performed in ImageJ using genexyz Polylux tool plugin from Resolve BioSciences.

##### Cell segmentation using Cellpose

Cellpose^51^ (v.2.0.4) was used to segment nuclei from the DAPI images with the pretrained nuclei model and flow_treshold 0.5, cellprob_threshold −0.2. The nuclear segments were then expanded by 10 pixels (1.38 μm) using the ‘expand_labels’ function implemented in scikit-image and transcripts were subsequently assigned to the expanded segments. Segments larger than 4 median absolute deviation (MAD) plus the median segment area were removed from the analysis. During clustering and cell type annotation low quality clusters of cells were removed which could not be properly annotated.

##### Normalization, clustering and annotation

Count matrices of segmented cells were normalized and scaled using the ‘SCTransform’ function of Seurat and then 10 PCs were used for clustering in UMAP space. Cell clusters were then annotated using marker genes from sphere-seq analysis; cells that could not be properly annotated were removed from the analysis. Annotation was then projected on cells as an overlay on Molecular Cartography DAPI images in ImageJ.

##### Feature Area Integration

Signals for landmark genes defining central (*Cyp2e1*), portal (*Cyp2f2*) and metastatic areas (*Gpx2*) were used to visualize different areas, then areas were manually drawn and their x and y coordinates were exported. Coordinates from different areas of the image were then matched with x and y coordinates of cell segmented count data to annotate single cells by their area of origin.

##### Differential gene expression (DGE) analysis between two groups

DGE analysis was done as described for sphere-seq analysis. But instead of using the sum of single cell counts across spheres, counts were summed across spatial feature areas.

##### Analysis of differences in cell type abundance between two groups

Cell type proportions were analyzed as described for sphere-seq analysis.

##### Colocalization analysis

A spatial neighborhood graph was constructed based on the Euclidean distance in 2D space of the centroids of the segmented areas. In this graph, vertices represent the cells which are connected by an edge if the distance is smaller than 10 μm. To construct the graph, we utilized a kd-tree based nearest neighbor search in a pre-defined radius of 10 μm as implemented in the R function ‘nn2’ (RANN v.2.6.1,searchtype=‘radius’) with a sufficiently large k (k=41). This approach runs in O(M logM) time and avoids computation of the distance matrix for thousands of objects. The resulting adjacency matrix was then used to construct a graph using the igraph^52^ package (v.1.3.4). From this graph the number of edges between cell types was computed in each region of interest (ROI) and divided by the sum of the number of cells for each cell type pair to normalize for total cell numbers in the annotated regions. For each slide the difference in the normalized number of edges between the two groups of ROIs, e.g. proximal versus distal, was subsequently computed. This value was compared to an empirical null distribution derived from randomly permuting the labels of the vertices (m=1000) per slide. This approach takes tissue composition and spatial structure into account and allows the computation of P values as P=(b+1)/(m+1) where b is the number of times the permutation produced a more extreme number of edges between two cell types than observed and m the total number of permutations^53^. This was done for each slide and possible cell-cell interactions to derive a score that represents the fraction of images in which a specific interaction was significant, with the sign representing co-localisation or avoidance; visualization was adopted from^54^.

## Data availability

The datasets generated in this study are deposited on Gene Expression Omnibus (GEO) with accessory number GSE216189 and zenodo.org with accessory number 10.5281/zenodo.7197153.

## Code availability

The code generated in this study is available at: https://github.com/Moors-Code/Sphere-sequencing

---

**Supplementary Fig. 1:**
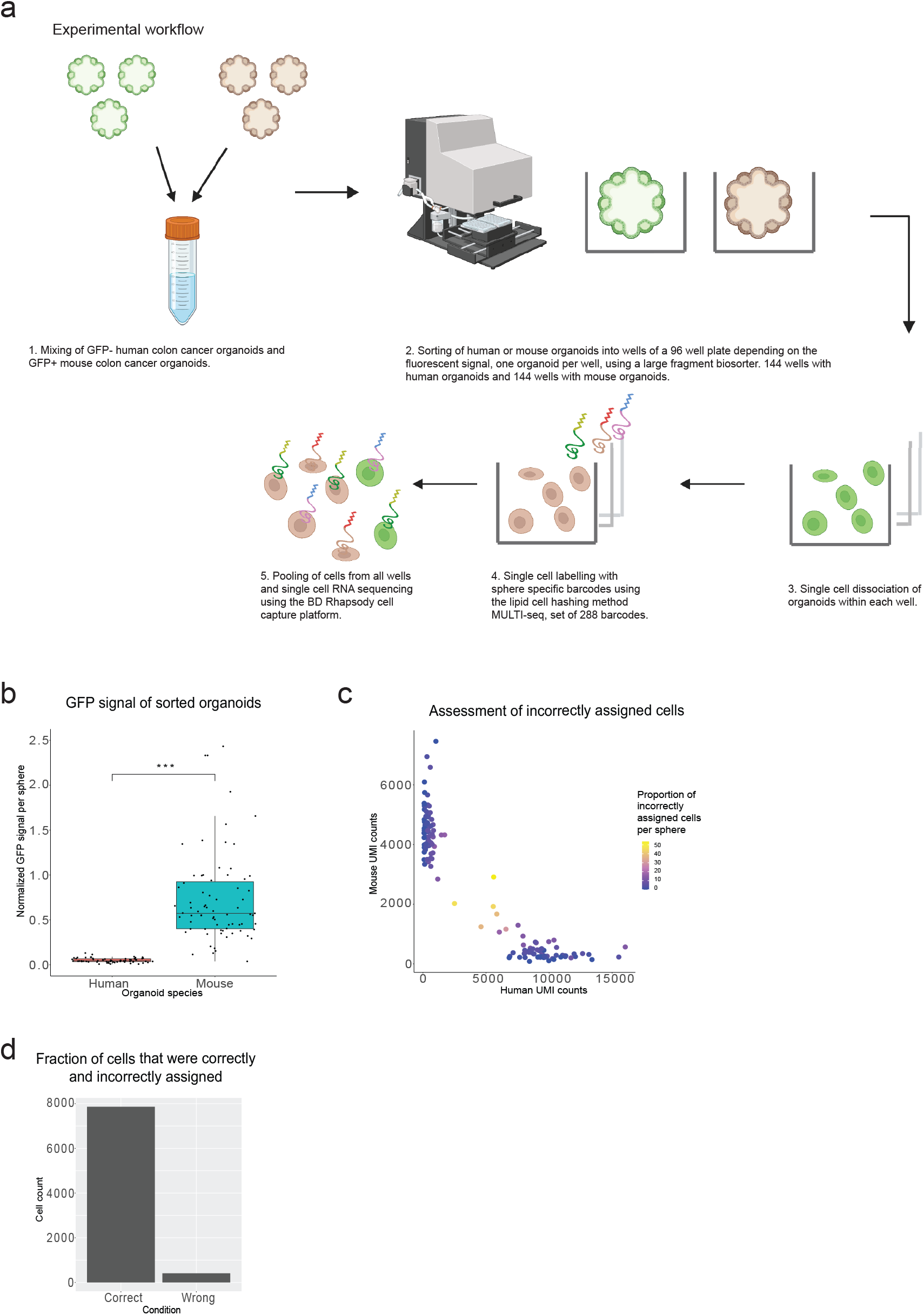
Index-sorting of mixed-species organoids demonstrates the high accuracy of sphere-sequencing. **a**, Schematic drawing of experimental workflow. **b**, Boxplot of normalized GFP signal acquired from sorted GFP-human and GFP+ murine organoids (n = 144 spheres each across 1 sample). Black dots represent individual spheres. P-value was calculated with a non-parametric wilcoxon signed-rank test (*** < 0.001). **c**, Scatter plot showing the fraction of incorrectly assigned cells. The x- and y-axis show the average number of unique molecular identifiers (UMI). Dots represent spheres and their color indicates the percentage of incorrectly assigned cells (human cells within mouse spheres and mouse cells within human spheres) (n = 139 spheres across 1 sample). **d**, Barplot showing the fraction of cells that were correctly and incorrectly assigned (n = 1 sample).

**Supplementary Fig. 2:**
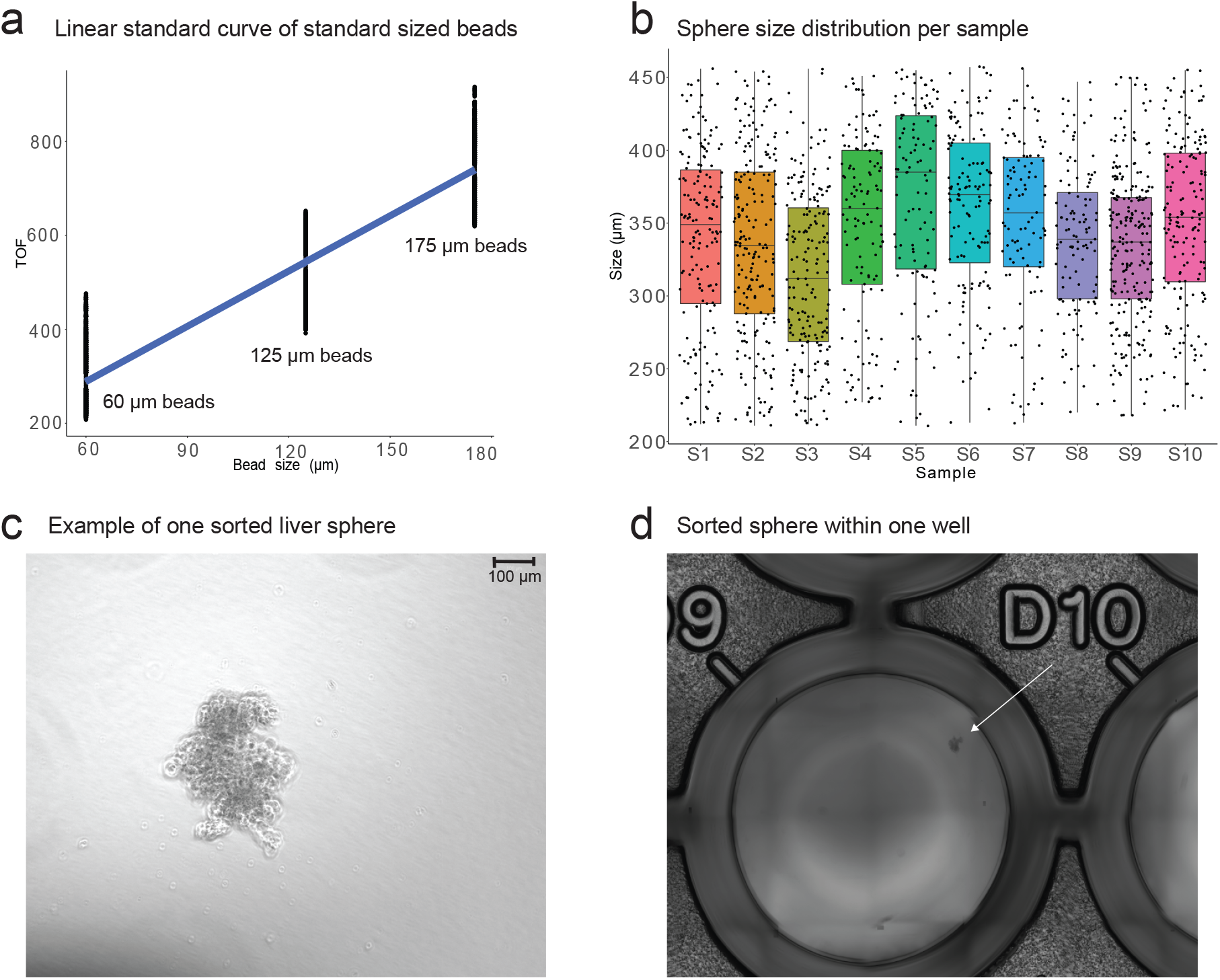
Sorting of spheres in a size dependent manner with a large fragment sorter. **a**, Linear standard curve generated by acquiring standard sized beads [60 μm (n= 982), 125 μm (n = 736) and 175 μm (n = 923)]. A linear model is fitted to calculate sphere-sizes. Black dots represent acquired measurements of individual beads. TOF: time of flight. **b**, Sphere-size distribution of sorted spheres per sample visualized in boxplots (n = 288 spheres per sample). Black dots indicate spheres. **c**, Brightfield microscopy image of a representative liver sphere that is around 300 μm in diameter, the scale bar is shown on the top right corner. **d**, Brightfield microscopy image of one well of a 96 well plate with a sorted liver sphere (white arrow).

**Supplementary Fig. 3:**
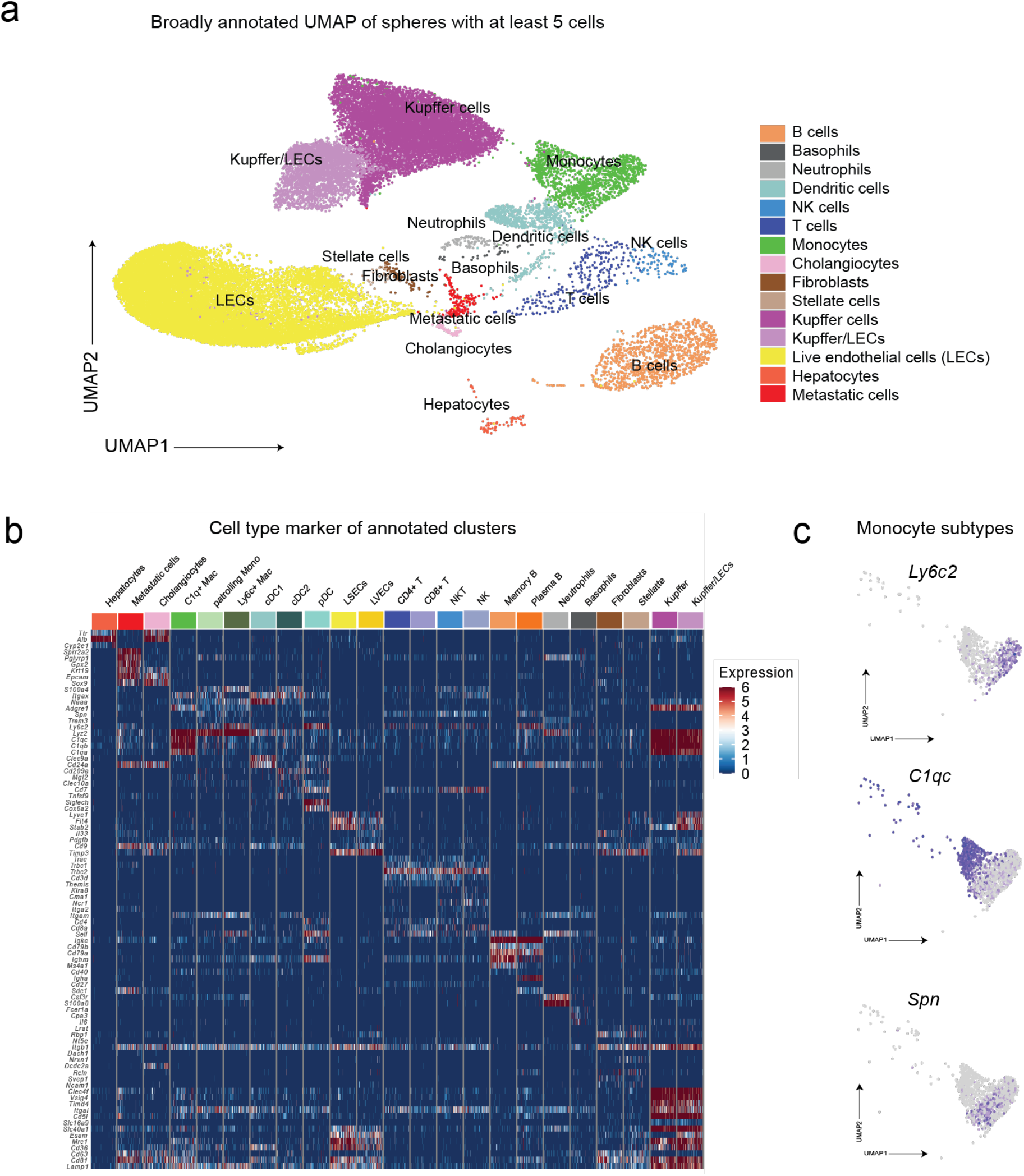
Single cell annotation of sphere-sequencing data. **a**, Uniform manifold approximation and projection (UMAP) visualization of integrated liver samples (n = 10: 9 injected with colon cancer organoids, 1 untreated). Cells are clustered, broadly annotated and coloured by their cell type. **b**, Heatmap showing marker genes used for cell type annotation. The columns represent cells and the rows represent genes. Expression levels per cell type cluster are highlighted in a gradient from blue to red. **c**, Feature plots showing expression of marker genes for different subsets in monocytes. Upper plot, *Ly6c2* indicating *Ly6c+* macrophages; middle plot, *C1qc* indicating *C1q+* macrophages; lower plot, *Spn* indicating patrolling monocytes.

**Supplementary Fig. 4:**
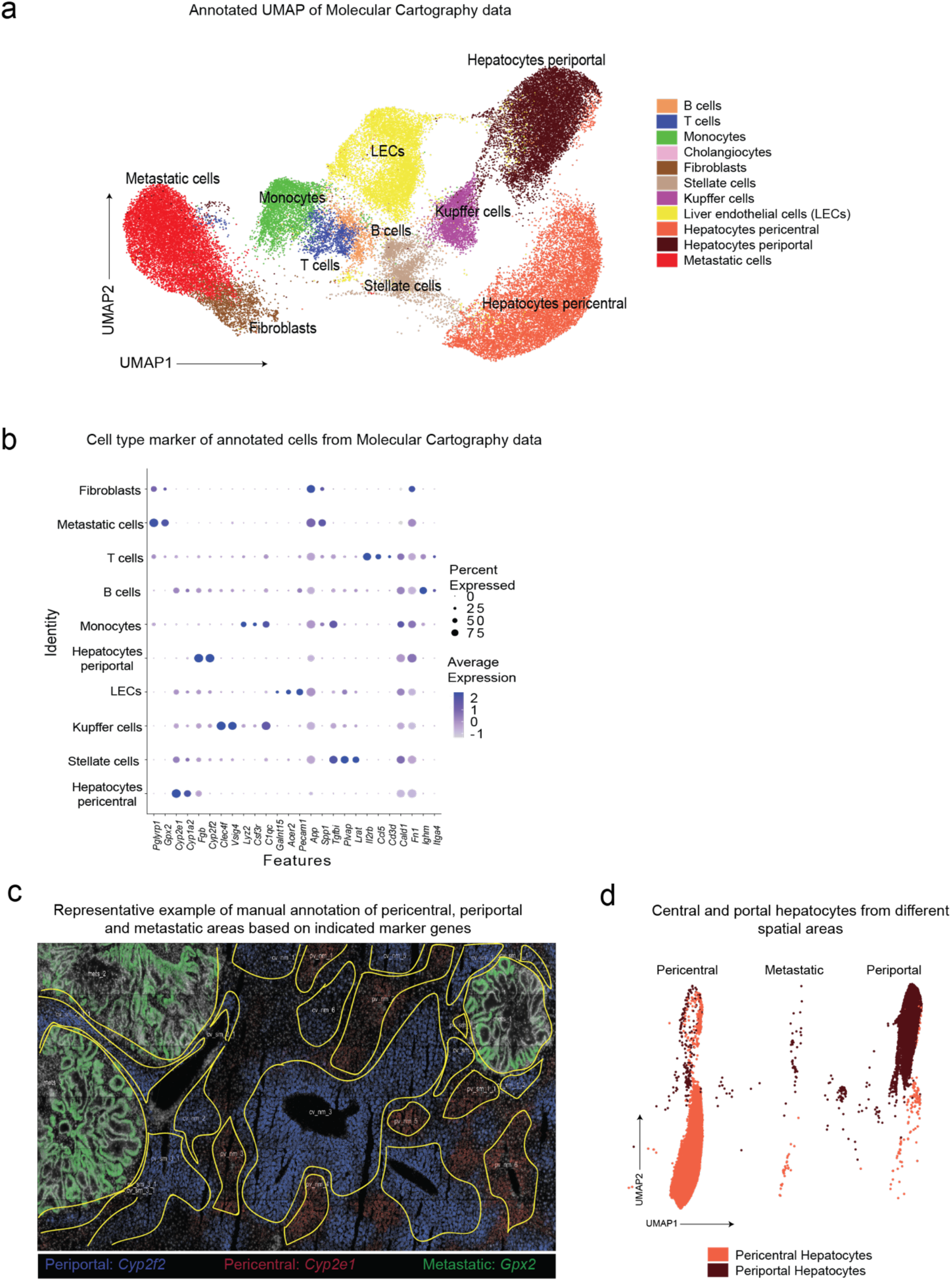
Single cell annotation of Molecular Cartography data. **a**, Uniform manifold approximation and projection (UMAP) visualization of single cells after cell segmentation (n = 4 samples). Cells are clustered, annotated and coloured by their cell type. **b**, Dotplot showing marker genes used for cell type annotation. **c**, Representative example of Molecular Cartography of indicated genes for periportal (*Cyp2f2*, blue), pericentral (*Cyp2e1*, red) and metastatic (*Gpx2*, green) areas. Yellow lines denote manually drawn areas grouping cells into different spatial zones. **d**, UMAP visualization of hepatocytes from different spatial zones. Cells are grouped and annotated based on their hepatocyte affiliation; hepatocytes found periportal in dark red or pericentral in light red.

**Supplementary Fig. 5:**
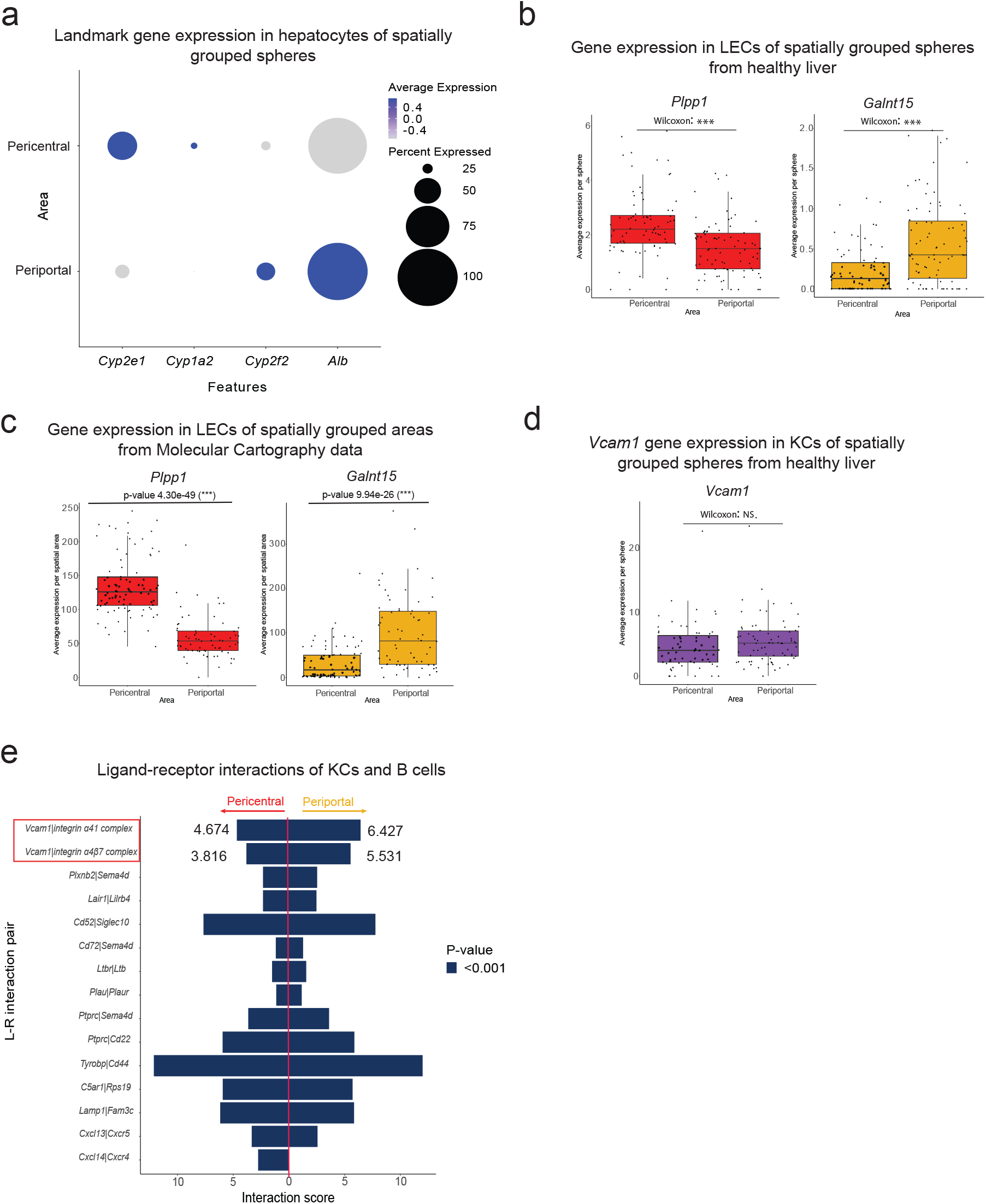
Sphere-sequencing application to mouse liver zonation. **a**, Dotplot showing hepatocyte landmark genes of grouped spheres in pericentral zones: *Cyp2e1, Cyp1a2* or periportal zones: *Cyp2f2, Alb* (pericentral n= 46, periportal n=31 spheres across 10 samples). **b**, Boxplots of zonated genes in LECs of spatially ordered spheres in healthy liver (pericentral n = 76, periportal n= 83 spheres across 1 sample). Black dots denote spheres. **c**, Boxplots of *Plpp1* (left) and *Galnt15* (right) in LECs of spatially grouped areas of Molecular Cartography data (pericentral n= 89; periportal n=66 areas across 4 samples). Black dots indicate individual spatial areas. **d**, Boxplots of *Vcam1* in Kupffer cells (KCs) of spatially ordered spheres in healthy liver (pericentral n = 67; periportal n= 74 across 1 sample). Black dots indicate individual spheres. **e**, Predicted ligand-receptor (L-R) interactions between monocytes and B cells in distal or proximal areas based on sphere-seq data (n=3 samples). Interaction scores were calculated by CellPhoneDB, which uses a permutation test to generate p-values indicating significantly enriched L-R interactions. Interactions referenced in the main text are highlighted with red squares and numbers indicate interaction scores. For **b** and **d** a non-parametric wilcoxon signed-rank test was applied (*** < 0.001, not significant > 0.05). For **c** a negative binomial generalized log-linear model was used for statistical testing and p-values (benjamini-hochberg adjusted) <0.05 were considered significant (***<0.001, **<0.005, *<0.05).

**Supplementary Fig. 6:**
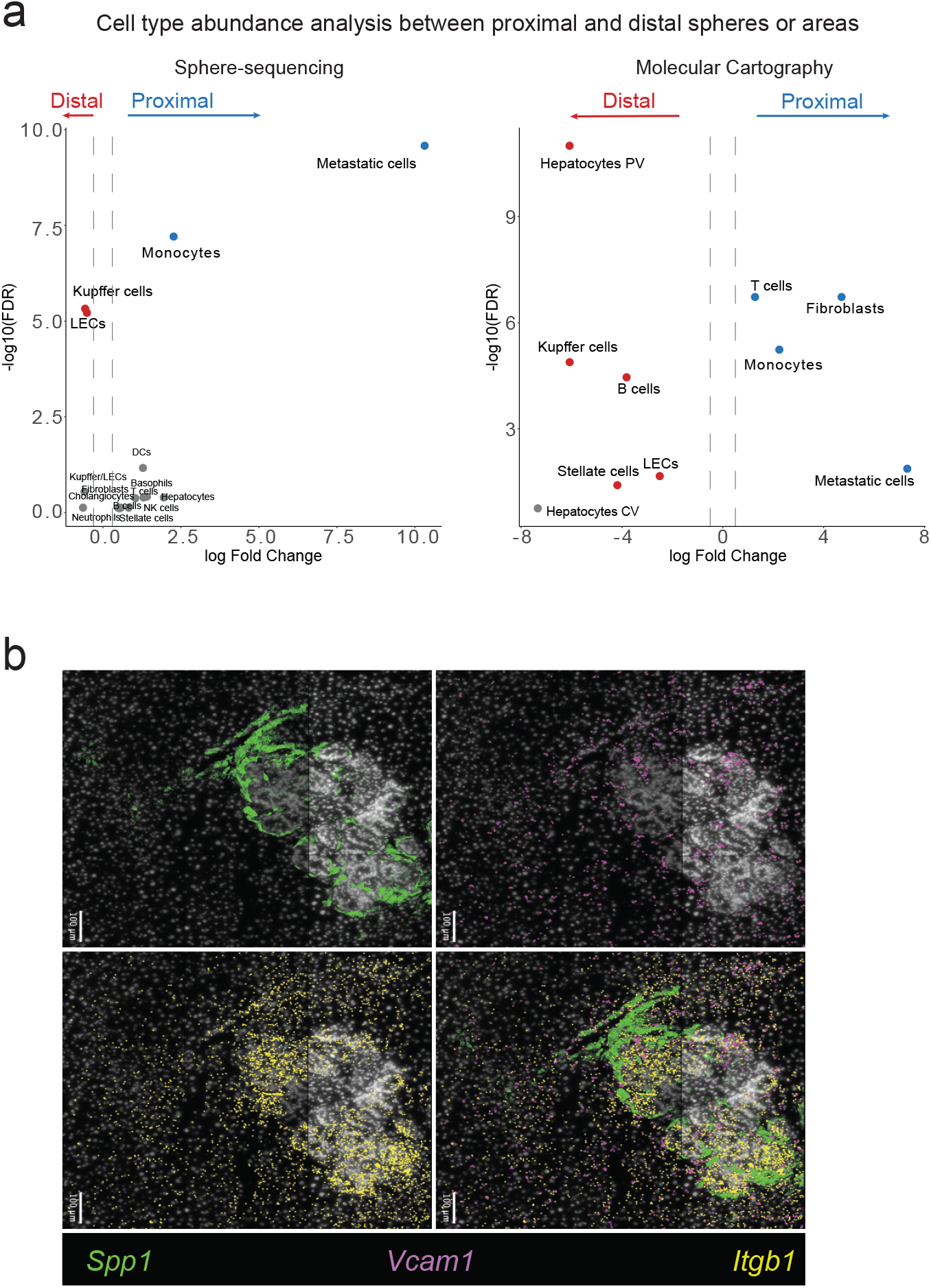
Sphere-sequencing application to mouse metastatic liver. **a**, Differential abundance analysis (DAA) of cell types between areas distal or proximal to metastases in sphere-seq data (left, n = 3 samples) and Molecular Cartography (right, n= distal: 71 areas, proximal: 11 across 2 samples). Colored dots represent significantly higher proportions of cell types; red, higher in distal; blue, higher in proximal. A negative binomial generalized log-linear model was used for statistical testing and p-values (benjamini-hochberg adjusted) <0.05 were considered significant. **b**, Representative Molecular Cartography of *Spp1* (green), *Vcam1* (purple) and *Itgb1* (yellow) shown as an overlay with DAPI signal (white).

**Supplementary Fig. 7:**
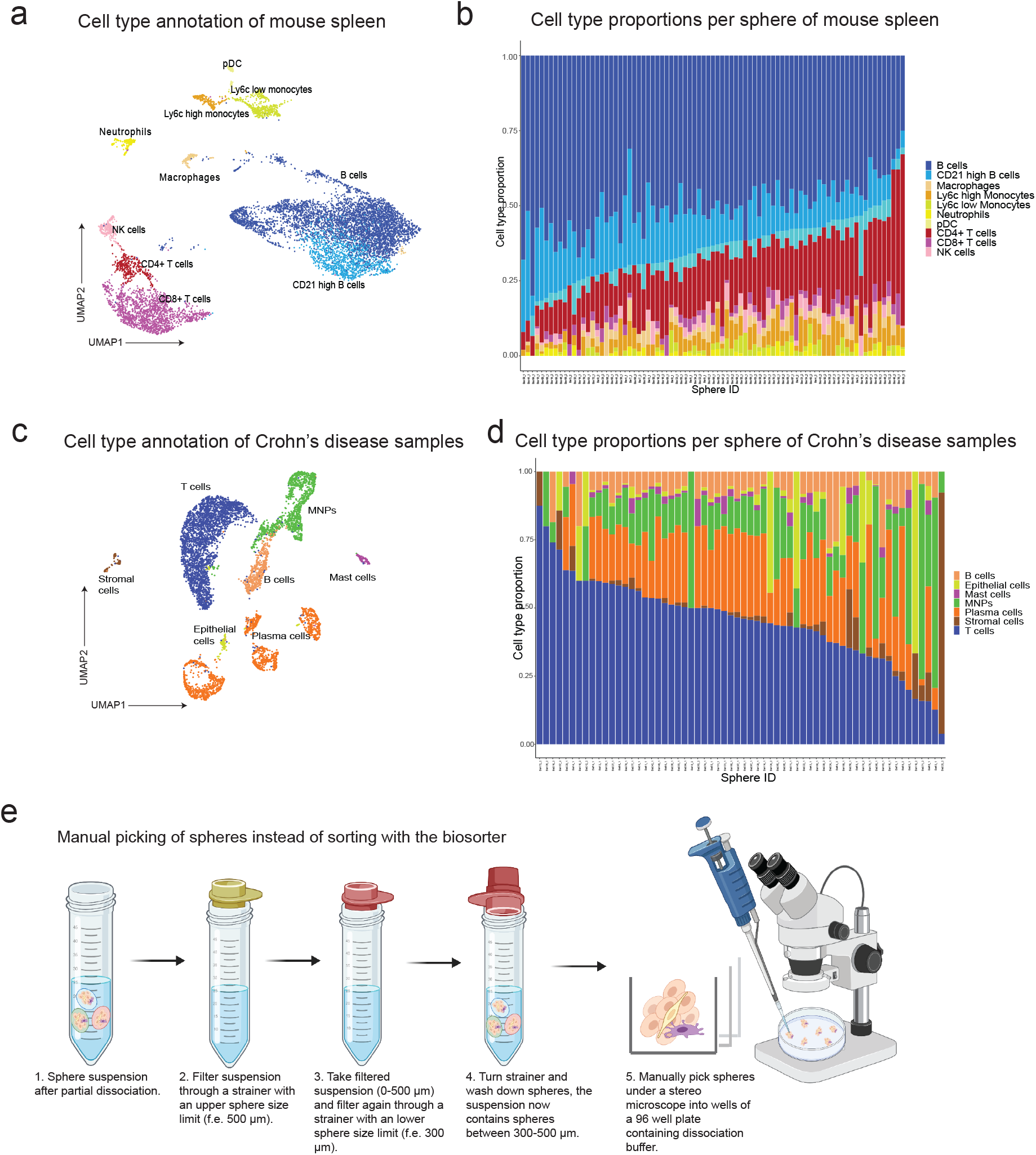
Proof-of-concept of applying sphere-sequencing to other tissues and species. **a**, Uniform manifold approximation and projection (UMAP) visualization of integrated sphere-seq data of mouse wild type spleen (n = 2); only spheres with at least 5 cells are considered. Cells are clustered, annotated and coloured according to their cell type. **b**, Barplot showing cell type proportions per sphere (n = 83 spheres across 2 samples). **c**,**d**, Same visualization like in a and b, but for Crohn’s disease biopsy samples (n = 62 spheres across 2 samples). MNP: mononuclear phagocyte. **e**, Schematic drawing of size selection of spheres for manual picking instead of using the large fragment sorter.

**Supplementary Fig. 8:**
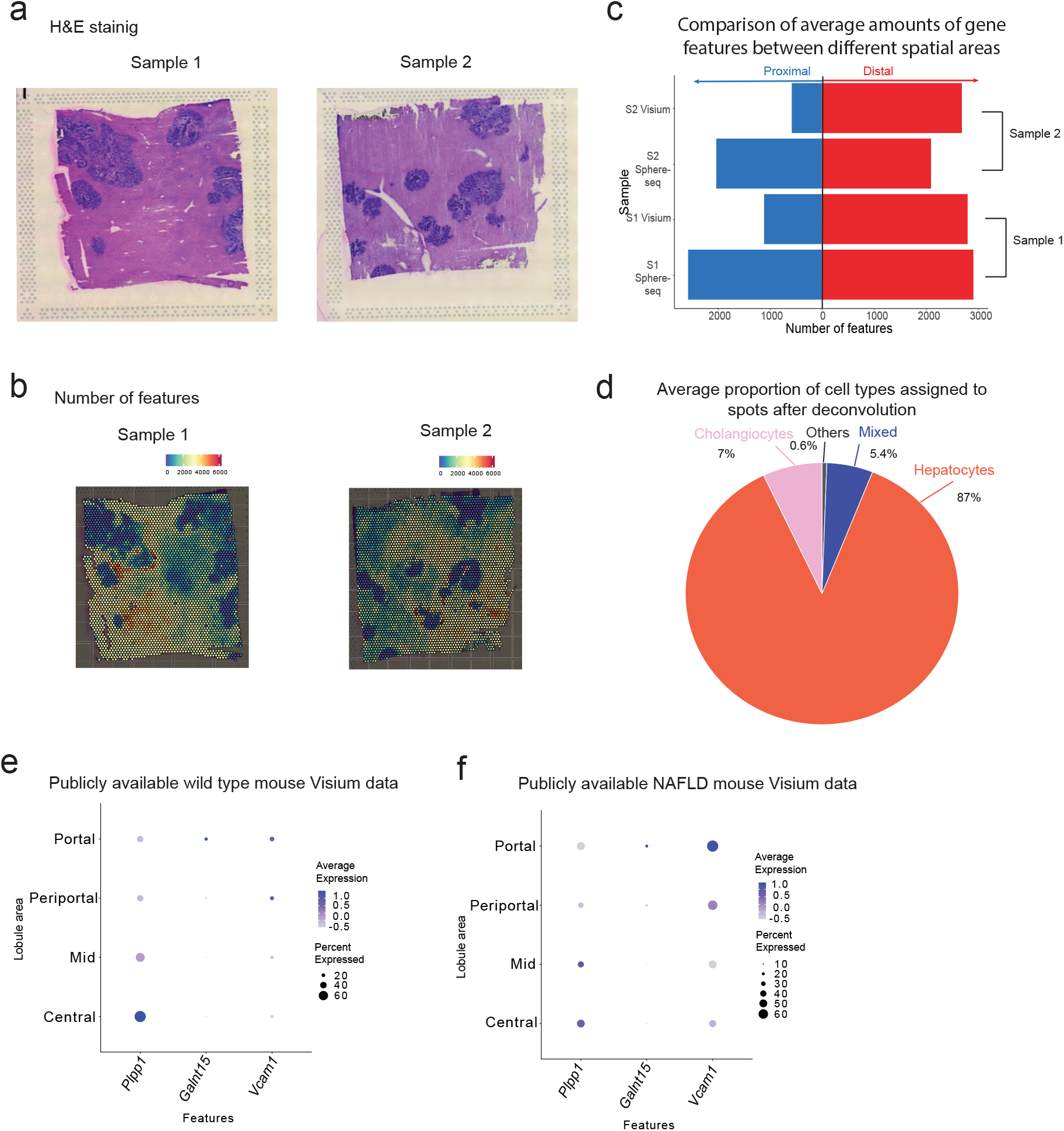
Comparison of sphere-sequencing with Visium. **a**, Hematoxylin and eosin (H&E) staining of mouse metastatic livers (n = 2). Metastatic areas are indicated by dark purple stains. **b**, Number of gene features per spot projected on H&E stainings from a. **c**, Boxplot comparing the number of features from different spatial areas of sphere-seq data to Visium (n = 2 each). The same two samples were used for sphere-seq and Visium, indicated by brackets. **d**, Piechart indicating the average proportions of cell types after spot deconvolution of integrated mouse metastatic liver samples (n=2). Spots were assigned to one single cell type if at least 75% of transcripts could be assigned to that cell type; other spots were annotated as mixed. **e**, Dotplot showing average expression of zonated genes *Plpp1, Galnt15*, and *Vcam1* in publicly available Visium data from healthy mouse livers^10^. **f**, Dotplot like in e from publicly available non-alcoholic fatty liver disease (NAFLD) mouse Visium data^10^.

